# Self-organized emergence of hyaline cartilage in hiPSC-derived multi-tissue organoids

**DOI:** 10.1101/2021.09.21.461213

**Authors:** Manci Li, Juan E. Abrahante, Amanda Vegoe, Yi Wen Chai, Beth Lindborg, Ferenc Toth, Peter A. Larsen, Timothy D. O’Brien

## Abstract

Despite holding great therapeutic potential, existing protocols for *in vitro* chondrogenesis and hyaline cartilage production from human induced pluripotent stem cells (hiPSC) are laborious and complex with unclear long-term consequences. Here, we developed a simple xeno- and feeder-free protocol for human hyaline cartilage production *in vitro* using hydrogel-cultured multi-tissue organoids (MTOs). We investigate gene regulatory networks during spontaneous hiPSC-MTO differentiation using RNA sequencing and bioinformatic analyses. We find the interplays between BMPs and neural FGF pathways are associated with the phenotype transition of MTOs. We recognize TGF-beta/BMP and Wnt signaling likely contribute to the long-term maintenance of MTO cartilage growth and further adoption of articular cartilage development. By comparing the MTO transcriptome with human lower limb chondrocytes, we observe that the expression of chondrocyte-specific genes in MTO shows a strong correlation with fetal lower limb chondrocytes. Collectively, our findings describe the self-organized emergence of hyaline cartilage in MTO, its associated molecular pathways, and its spontaneous adoption of articular cartilage development trajectory.

## Introduction

Cartilage tears and osteochondral defects involving the articular cartilage are common injuries that may lead to disabling osteoarthritis and subsequent joint replacement. Injured articular cartilage has a limited capacity for self-regeneration, and currently, available treatment options for symptomatic cartilage lesions are limited. Surgical repair methods such as osteochondral auto/allograft transplantation and bone marrow stimulation via microfracture all have their limitations including the scarcity of suitable grafts or their inability to yield repair tissues composed of hyaline-vs. fibrocartilage. To address these shortcomings, increasing efforts have recently focused on implementing cell-based methods for the treatment of articular cartilage injuries. First and second-generation autologous chondrocyte implantation methods have been shown to yield good patient outcomes, but the morbidity and cost associated with the surgical harvest of autologous chondrocytes for *in vitro* expansion remain critical drawbacks^1^.

In order to eliminate the pitfalls associated with the surgical harvest of autologous chondrocytes, studies have lately concentrated on *in vitro* chondrogenesis and hyaline cartilage production from human stem cells^2,3^. Specifically, human induced pluripotent stem cells (hiPSC) have been shown to have the potential to differentiate into chondrocytes; however, clinical translation of hiPSC derived chondrocytes still faces several challenges. The common approach for generating hiPSC-derived chondrocytes has been a 2D-3D sequential culture where hiPSC-derived mesodermal cells are cultured in monolayer before 3D—such as pellet^4-7^ and suspension^8,9^—cell culture. Implementing 3D culturing in the process has been shown to improve the quality of the cartilage generated^10^. Nevertheless, these existing step-wise protocols are labor-intensive and involve the use of fetal bovine serum (FBS) as well as manipulations of inductive and repressive signals for mesoderm specification in embryonic development^6^. Moreover, the long-term maintenance and consequences of chondrocytes in suspension and pellet cell cultures remain to be explored.

Organoids are 3D cultures that realize the self-organizing potential of stem cells^11,12^. Several transcriptome analyses have shown that organoids can recapitulate a variety of early developmental processes in human organs, such as brain^13-17^, retina^18^, kidney^19^, intestinal epithelium^20^, and trophoblast^21^. As chondrogenic tissue formation occurs at the highest levels during fetal/younger stages of life^22^, 3D organoid systems have a unique advantage in achieving *in vitro* chondrogenesis and even associated organization of extracellular matrices. *De novo* hyaline cartilage from bovine organoids has been recently reported and showed higher similarity to native cartilage^10^. Human clinical translation of organoid-derived hyaline cartilage and chondrocytes requires xenobiotic-free and serum-free culturing protocols^23,24^. Elucidation of key mesoderm formation pathways associated with cartilage production in organoids is needed to facilitate future cartilage production through organoid engineering.

Here, we report the spontaneous appearance—without the addition of external chemicals beyond those included in E8 medium—of hyaline cartilage in hiPSC-derived multi-tissue organoids (MTOs) using a modification of our previously reported method that employed Cell-Mate3D hydrogels and formed predominantly brain tissue^25^. To gain a better understanding of the molecular pathways in MTOs during *in vitro* cartilage production, we conducted RNA-Seq at weeks 8, 11, and 15 following MTOs induction. By comparing with existing RNA-seq data obtained from human lower limb chondrocytes at different life stages^6,26-28^, we demonstrate that the relevant gene expression in 15-week MTOs is strongly correlated with human fetal lower limb tissues. In summary, this study describes the spontaneous emergence of human hyaline cartilage from hiPSC-derived MTOs cultured with a xeno- and feeder-protocol and its associated molecular pathways.

## Results

### Culturing hiPSC to cartilage producing MTO

We have previously reported the generation of cerebral organoids (CO) from human pluripotent stem cells using a chemically defined hydrogel material (Cell-Mate3D) and culture medium (E8) and characterized their composition for up to 28 days from induction by which time they were approximately 3mm in diameter^25^. However, the culture time span of these organoids was limited due to central necrosis of the organoids when they reached 2-3mm in diameter, presumably due to hypoxia. To address this limitation, we adapted our methods to use a bioreactor system that has a gas-permeable bottom (GREX100) thereby allowing oxygen diffusion from the bottom as well as the top of the culture medium interfaces. Using this system along with continued use of only E8 medium, we were able to routinely culture organoids for months, following some for up to 30 weeks (week 30). A further simplification of our originally reported method involved the elimination of the use of the chitosan component of the Cell-Mate3D μGel. As early as week 6, we observed that, although the cerebral phenotype of the organoids was still prominent, cartilage-like tissues started to emerge—organoids with multiple tissue types are hereon referred to as multi-tissue organoids (MTOs) (Fig. S1). Notably, despite their apparent reduction in the relative proportion of total MTO tissue, neural tissues persisted in MTOs throughout the characterized period of development (Fig. S1). Cartilage, which formed centrally in the MTOs, was easily recognizable through histology due to its distinct morphology and characteristic histochemical and immunohistochemical features. The main types of cartilage—articular, hypertrophic, elastic, and fibrous cartilage—can be distinguished by the structure and composition of their extracellular matrix (ECM). Articular cartilage, for example, has a hyaline rather than fibrous morphology (fibrocartilage), and contains predominantly type II collagen, little or no type I collagen (fibrocartilage) or type X collagen (hypertrophic cartilage), and no elastic fibers. By week 8, hyaline cartilage became very distinguishable in MTOs, as indicated by the development of characteristic, abundant, homogenous pale basophilic extracellular matrix (ECM) in H&E stained sections (Fig. 1a) and prominent Alcian blue staining of proteoglycan/hyaluronic acid components typical of cartilage ECM^32^ (Fig. 1b). Immunohistochemistry of MTO cartilage showed increasing amounts of type II collagen over time as the cartilage matured and showed extensive expression of aggrecan at all time points (Fig. 1c-d). Type VI collagen was also extensively expressed in MTO cartilage while Type I collagen showed expression in some sites at the periphery of MTO cartilage, and Type X collagen generally showed no immunoreactivity above background (Fig. S2). We noted that the hyaline cartilage phenotype was stably maintained from week 8 to week 30 with prominent growth in size (Fig. 1a-d). To further assess this, we performed histomorphometric measurements on MTO histologic sections using aggrecan area fraction as a global indicator of developing and mature cartilage composition of the MTOs. At week 8 (n=7 biological replicates) the mean aggrecan area fraction was 17.7% (± 7.2) and at week 11 (n=5) was 57.8 (± 15.1) with unpaired t-test showing a significant increase from week 8 to 11 (p=0.001) (Fig. 1e).

**Fig. 1.**
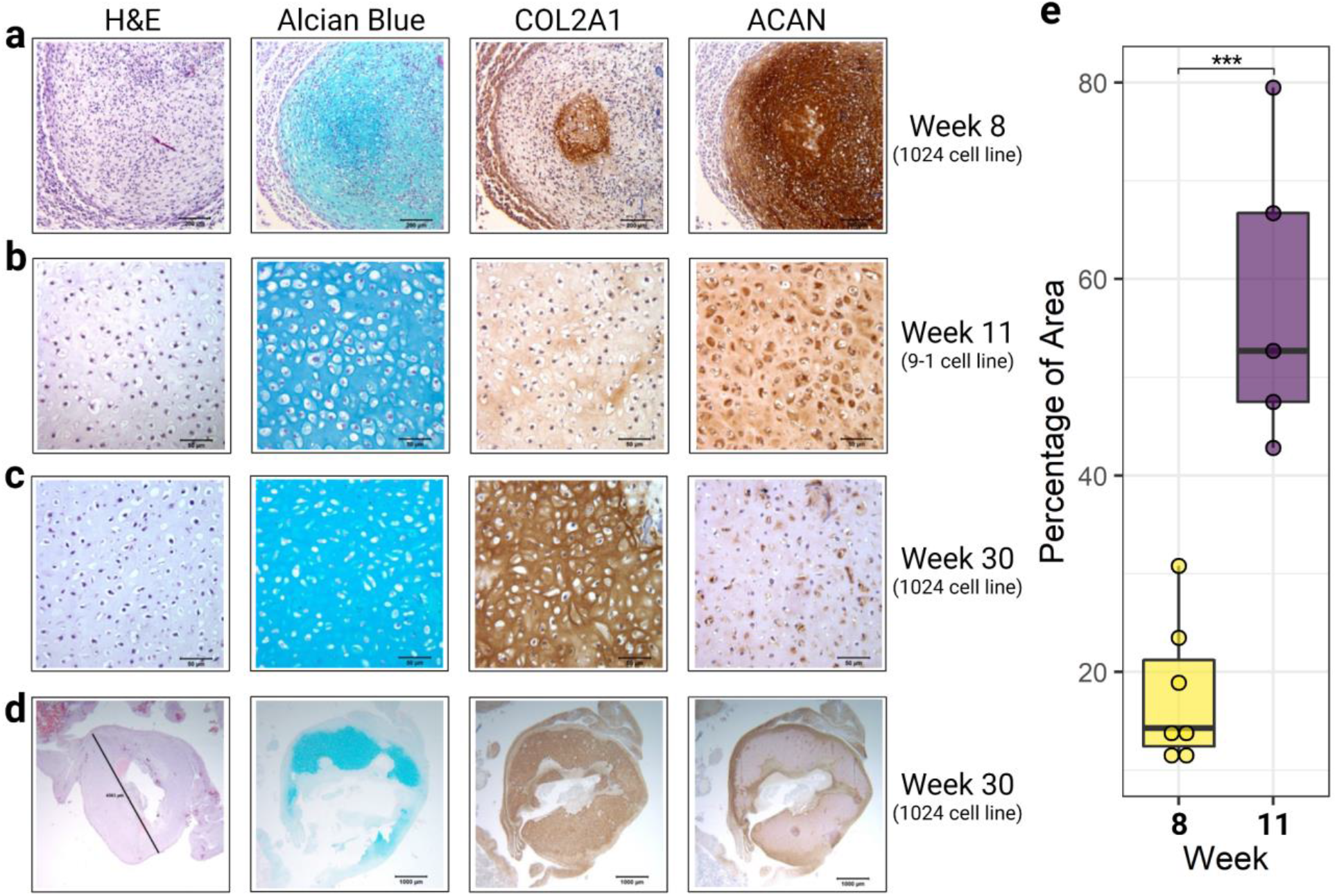
Histology of 1024 cell line multi-tissue organoids (MTOs) at 8 and 30 weeks, and 9-1 cell line^25^ MTOs at 11 weeks. **a.** MTOs at 8 weeks shows a developing cartilage nodule with diffuse Alcian blue staining (blue) of the early cartilaginous matrix, diffuse labeling for aggrecan (ACAN, brown) in cells and matrix, and type II collagen labeling (COL2A1, brown) in the cartilaginous matrix of a central region of maturing cartilage. Size bars = 200 μm. **b.** MTOs of the 9-1 cell line at 11 weeks show maturing cartilage with increased Alcian blue positive cartilaginous matrix separating chondrocytes, moderate diffuse staining for type II collagen, and diffuse aggrecan labeling. Size bars = 50 μm. **c.** 1024 MTOs after 30 weeks in culture show further maturation to hyaline cartilage morphology with chondrocytes surrounded by an abundant matrix with diffuse Alcian blue and type II collagen staining, and pericellular aggrecan staining. Size bars = 50 μm. **d.** Low magnification views of 1024 30-week MTOs shown in **c.** demonstrating the large size attained by some chondrogenic nodules. Measure bar on H&E panel = 4,363 μm (4.363 mm). Size bars = 1000 μm. **e.** Histomorphometric measurements on 1024 MTOs histologic sections using aggrecan area fraction (percentage) in MTOs at week 8 and 11 in 1024 cell line (***, p=0.001) (Fig. 1e).

### Global transcriptome reveals signatures of mesoderm formation in MTOs

To understand the transcriptome change underlying the phenotypic development of cartilage in MTOs, we conducted bulk RNA-seq on 1024-derived MTOs at weeks 8, 11, and 15, which covered the time span during which our histologic analysis showed emergence, expansion, and maturation of the MTOs cartilage. Principal component analysis (PCA) of global RNA expression data in MTOs showed distinct clustering corresponding to the time of collection (Fig. 2a). We conducted differential gene expression and gene ontology (GO) enrichment of differentially expressed genes for all three comparisons (Fig. S4; Fig. 2). We were especially interested in the altered expression of several cartilage marker genes at week 15 in comparison to week 8. Likely due to the multi-tissue nature of MTOs, although COL2A1 content increased in MTOs cartilage (Fig. 1c), it showed decreased expression in bulk RNA-seq. Other cartilage markers, however, such as *ACAN, CD44, COMP, PRG4*, and *SNAI1*, displayed significantly increased expression (Fig. 2b; Table S1-2). We note that although transcript levels for collagen type I/X increased, the expression of another hypertrophic marker, *IHH*, decreased, suggesting that further analyses into cartilage development pathways were needed. Additionally, type I collagen is the most abundant collagen and its expression is not limited to cartilage ^29^; therefore increased levels of type I collagen transcripts in MTOs do not necessarily reflect an increase of type I collagen in the composition of MTOs cartilage. Interestingly, we observed consistent downregulation of neural processes, such as synapse organization and axonogenesis, and upregulation of mesodermal processes, for example, extracellular matrix organization, connective tissue development, and cartilage development (Fig. 2c-d; Fig. S4; Table S3-4).

**Fig. 2.**
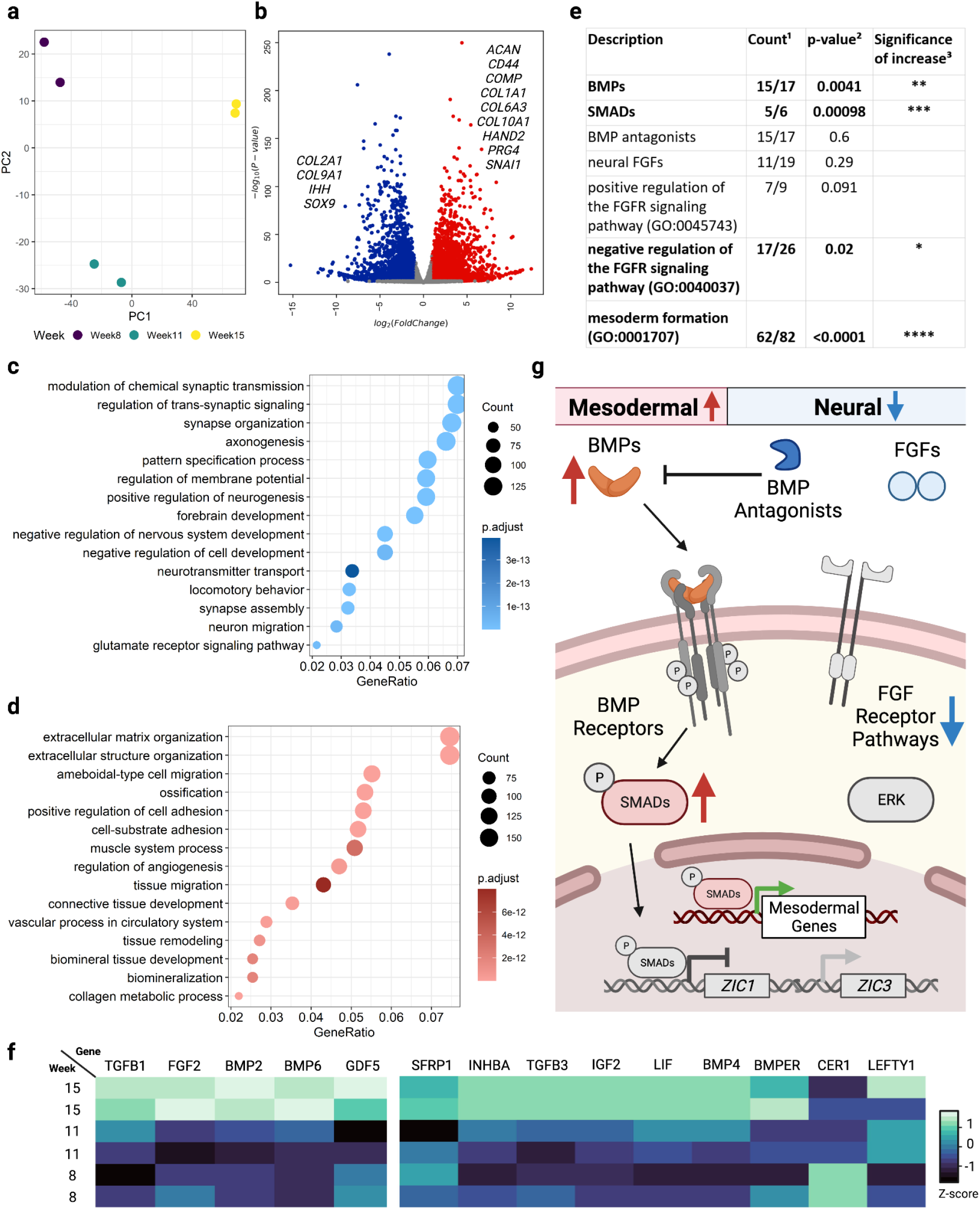
Cartilage development in MTOs is associated with favored BMP signaling pathways and increased mesodermal gene expression. **a.** Principal component analysis (PCA) of MTO global transcriptomes at weeks 8, 11, and 15. **b.** Differential gene expression analysis showing differentially expressed cartilage marker genes between week 8 and week 15. **c.** Ontology enrichment of differentially decreased genes between week 8 and week 15. **d.** Ontology enrichment of differentially increased genes between week 8 and week 15. **e.** Table displaying the comparison and statistical results of the overall expression of grouped genes between week 8 and week 15. ^1^ number of genes expressed/genes associated with the GO term (retrieved by biomaRt v2.46.3); ^2^ one-tailed Wilcoxon test comparing normalized and log-transformed transcript counts between week 8 and week 15; ^3^ *, p-value<0.05; **, p-value <0.01, ***, p-value <0.001, ****, p-value <0.0001. **f.** Expression of genes in MTOs at weeks 8, 11, and 15 encoding molecules commonly used for inducing chondrogenesis *in vitro*. **g.** Graphic summary (non-grey objects) of findings and relevant pathways.

Interplays between bone morphogenetic protein (BMP) and fibroblast growth factor (FGF) signaling play an important and highly conserved role in neural induction and mesoderm patterning, chondrocyte differentiation, and proliferation, and endochondral ossification during embryonic development studied almost exclusively in other species^30-33^. We observed a gradual increase in cartilage production from week 8 to week 15, therefore, we hypothesized that the global neural-to-mesodermal transition observed in human MTOs would be associated with significantly altered dynamics between BMP and FGF pathways. Specifically, we expected to see favored BMP pathways, which would entail 1) increased expression of components in the BMP signaling pathways, 2) reduced or unchanged levels of BMP antagonists, and 3) reduced or steady levels of FGF signaling. Pathways during mesoderm development processes are intertwined and expressed in a variety of tissues^30-32^; identifying and examining only a few expressed genes in one pathway may be biased in bulk-sequencing of MTOs. Therefore, we decided to examine the expression of genes comprehensively based on previous publications and GO terms and compared the overall expression of grouped genes between weeks 8 and 15. We observed that there was a significant increase in the overall expression of BMPs^34^ (p=0.0041) and their intracellular signaling transducers—SMADs^34^ (p=0.00098) (Fig. 2e; Fig. S4a-b). The overall expression of BMP antagonists^35^ remained unchanged (p=0.6) (Fig. 2e; Fig. S4c). As for FGF signaling pathways, we first examined the expression of neural FGFs^36^; which we found to be expressed at lower levels in comparison to other transcripts described previously although they did not change significantly (p=0.29) (Fig 2e; Fig. S4d). Further, we examined the dynamic expression of other components in the FGFR pathways and observed a significant increase (p=0.02) in negative regulation of the FGFR signaling pathway (GO:0040037) and a decrease—although not statistically significant (p=0.097)—in the expression of genes involved in the positive regulation of the FGFR signaling pathway (GO:0045743) (Fig 2e; Fig. S4e-f). This suggests that neural FGFs and their downstream pathways were suppressed, agreeing with the observed histological decrease in neural components. Finally, we investigated the expression dynamics of genes involved in mesoderm formation (GO:0001707) and we found a significantly increased (p<0.0001) expression of genes involved in this biological process (Fig 2e; Fig. S4g).

Because the MTOs spontaneously favored the phenotype of cartilage production without the addition of chemicals beyond the E8 medium, we wondered if MTOs intrinsically increased the expression of differentiation factors that are used to induce chondrogenesis. Gene products of *BMP2, BMP6, FGF2, GDF5*, and *TGFB1*—albeit still being debated—have been suggested to be indispensable for chondrogenesis^5,8^. Interestingly, expression of all of these genes—except *GDF5* (p=0.08)—significantly increased in MTOs spontaneously from week 8 to week 15 (Fig. 2f; Table S1). In addition, we examined other genes whose products were used for chondrogenic differentiation in other protocols^6,28^, including *TGFB3, INHBA, BMPER, BMP4, LIF, IGF2, LEFTY1, SFRP1*, and *CER1*. We found that all but *CER1* displayed increased patterns of expression (Fig. 2f; Table S1). This indicates that molecular mechanisms underlying spontaneous chondrogenesis in MTOs may resemble those observed in hiPSC-induced chondrogenic differentiation.

Overall, the global transcriptome of hiPSC-derived MTOs showed that increased gene expression in BMP signaling and mesoderm development, as well as decreased gene expressions in neural FGF signaling were associated with the increased cartilage production in MTOs from week 8 to week 15 (Fig. 2g).

### Distinct signaling pathways associated with increased articular cartilage development in MTOs

The establishment of both cartilage and bone formation is the result of chondrogenesis, which plays an essential role during the fetal development of the mammalian skeletal system. Hypertrophy of chondrocytes and deterioration of cartilage matrix precede endochondral ossification that leads to the formation and growth of long bones^37^. Gene expression for chondrogenesis and ossification overlap greatly due to the unified nature of cartilage and bone formation. As previously mentioned, we observed an increase in several cartilage marker genes as well as contradicting expression of *IHH* and *COL10A1* in the MTOs (Fig 1). Notably, we did not observe the deterioration, mineralization, or osteogenesis of the cartilage matrix even up to 30 weeks. Therefore, we probed further into the temporal expression patterns of genes crucial for cartilage development to see if we could identify signaling pathways that may contribute to the long-term maintenance of chondrocytes. To achieve this, we first investigated gene expression under the GO term cartilage development (GO:0051216) which includes both the positive and negative regulators of cartilage development^35^. We visualized the expression pattern of 125 (FDR<0.05) out of the 194 genes retrieved by biomaRt and found a clear separation of 42 and 83 genes with decreased and increased expression, respectively (Fig. 3a)^38^. In addition, despite the decrease of 42 genes, the overall gene expression levels increased significantly (p<0.0001) for cartilage development (Fig. 3a), agreeing with the expansion of cartilage in MTOs. We hypothesized that there would be distinct interactions of gene products between selected genes with increased and decreased expression. Therefore, we used STRING to construct two functional association networks consisting of proteins expressed by the described genes. Transcription factors and growth factors were the most connected molecules for both networks. Interestingly, we found that Wnt signaling molecules constituted the most prominent local network clusters in the network cluster organization for decreased gene products (Fig. 3b; Table 1; Table S5-6). Wnt signaling cascades have essential roles in the development and homeostasis of chondrogenesis and ossification^39^. In general, canonical Wnt cascades—including ROR2 and SFRP2 highlighted in the network—inhibits the early stages of chondrogenesis^38^. *WNT7A* overexpression blocked early chondrogenesis in chick limb model^40^. During endochondral ossification, Wnt signaling pathways promote chondrocyte hypertrophy^39^. SFRP2, WNT01B, and WNT7B are known positive regulators of ossification^35^ (Fig. 3a). Additionally, some non-Wnt positive regulators of ossification were also downregulated. Among the upregulated members, we observed clusterings centered on TGF-beta/BMP signaling pathways (Fig 3c; Table 1; Table S5-6). This is not surprising given TGF-beta/BMP signaling pathways predominantly promote all stages of chondrogenesis^41,42^. We also found a unique signature of increased downregulation of the canonical Wnt signaling pathway, agreeing with the association network formed by protein products of genes with decreased expression. In contrast to the previous network, we observed that several negative regulators of ossification mostly did not overlap with TGF-beta/BMP signaling pathways (Fig. 3c; Fig. S4). In summary, from week 8 to week 15, the cartilage in MTOs was likely stably growing due to continued increase in TGF-beta/BMP pathways promoting chondrogenesis and downregulation of Wnt signaling cascades that inhibit chondrocyte hypertrophy and ossification in maturing chondrocytes.

**Fig. 3.**
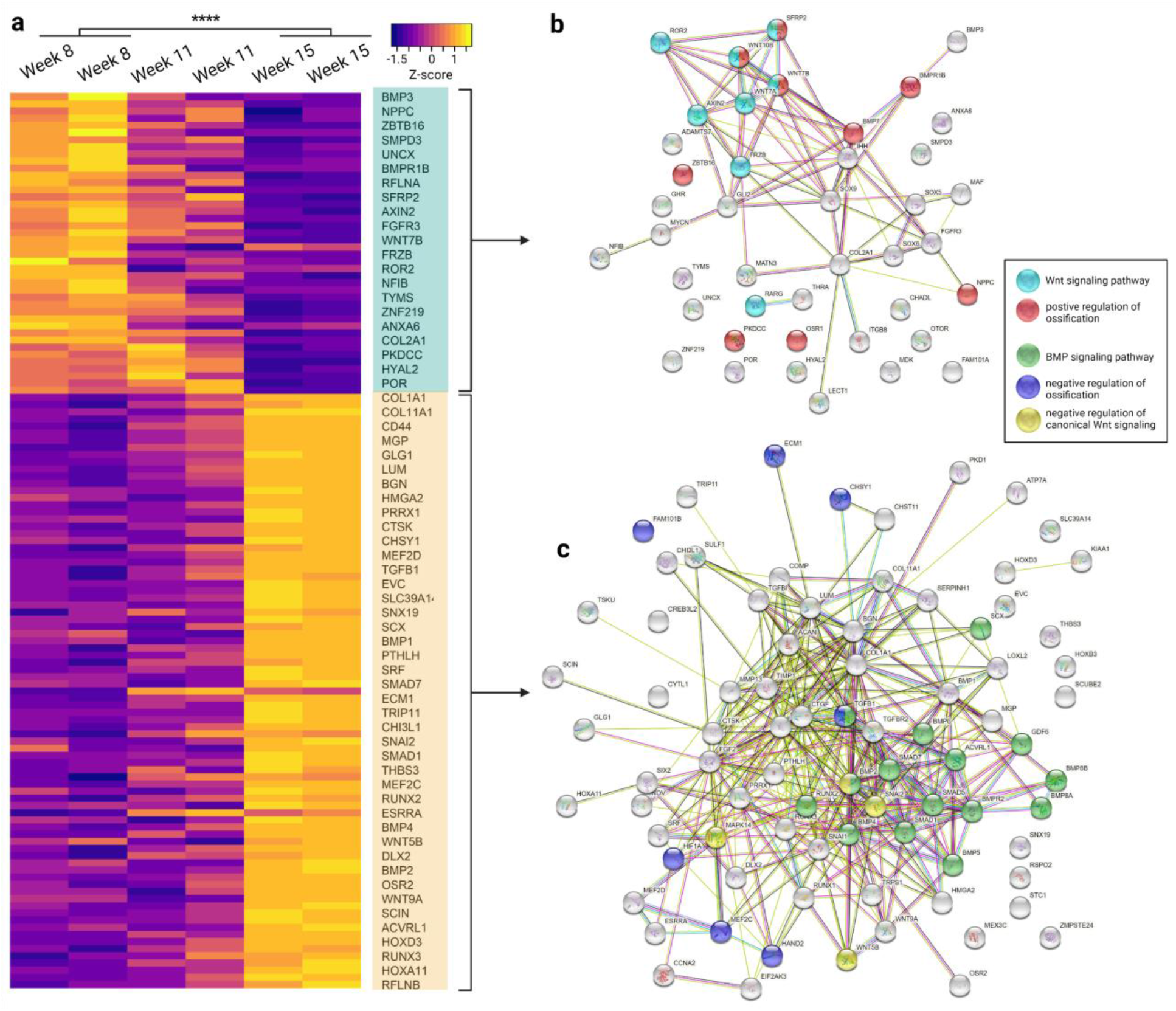
Cartilage development in MTOs is associated with distinct Wnt and TGF-beta/BMP signaling. **a.** 125 genes were selected (FDR<0.05) and their scaled temporal expression at weeks 8, 11, and 15 were plotted; genes were arranged by decreased and increased expression. Gene expression between week 8 and week 15 was compared. ****, p<0.0001. **b.** Functional association network of protein production expressed by genes with decreased expression. **c.** Functional association network of protein production expressed by genes with increased expression.

**Table 1.**
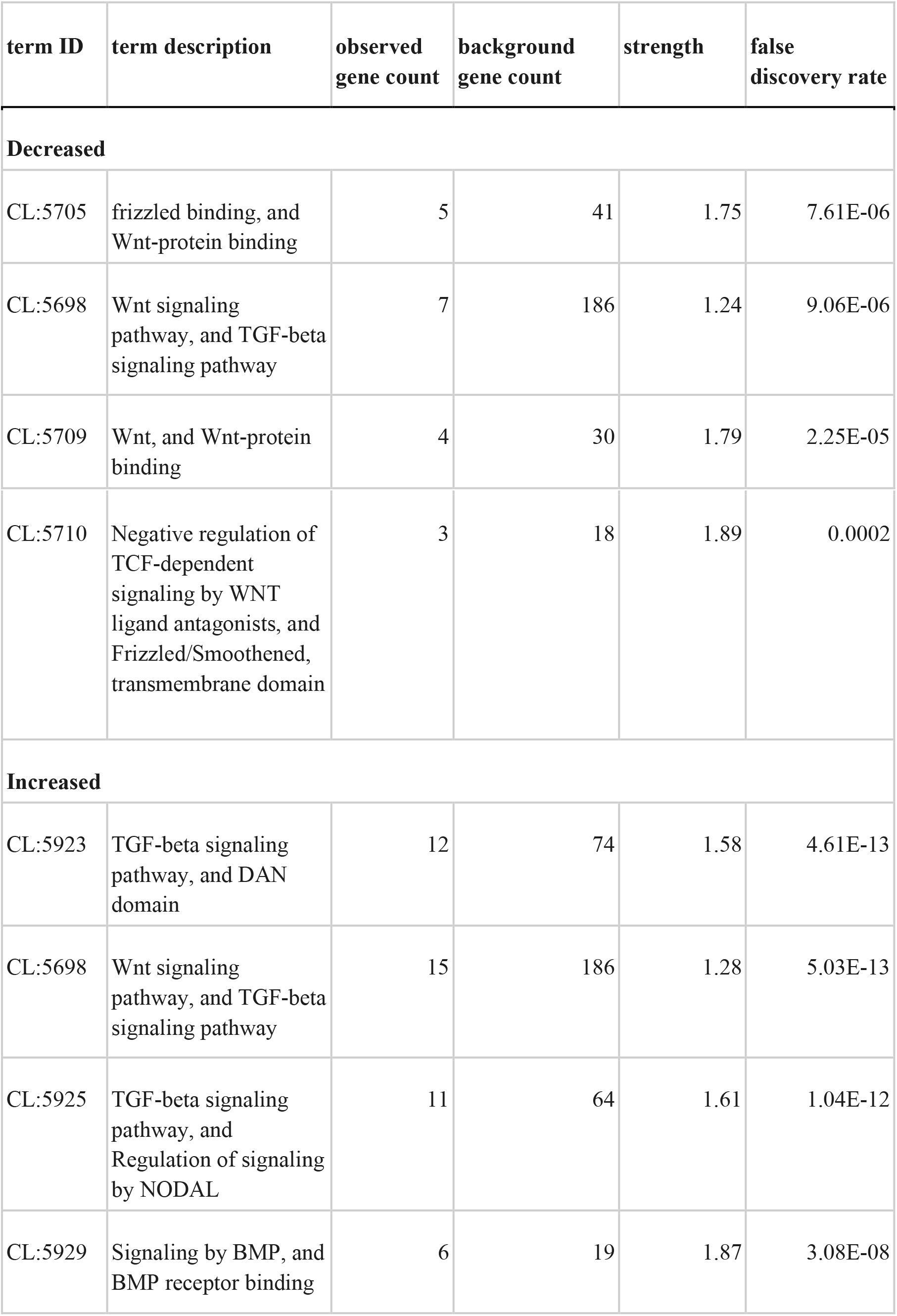
Top local network clusters (STRING)

In addition, we also compared the expression of genes for biological processes of interest as described above (Table 2; Table S7) using GO term gene lists retrieved by biomaRT from GO to reduce bias in gene selection^35^. Again, we found that genes in GO terms associated with promoting cartilage maturation, including positive regulation of cartilage development (GO:0061036), chondroblast differentiation (GO:0060591), and cartilage morphogenesis (GO:0060536), demonstrated significantly increased expression (Table 2; Table S7), while growth plate cartilage development gene set (GO:0003417) was not significantly increased (p=0.058). Moreover, the expression of genes associated with negative regulation of chondrocyte maturation and promotion of endochondral ossification did not show significant changes (Table 2; Table S7). As we observed the marginally insignificant increase of growth plate cartilage development (GO:0003417) marker gene expression (p=0.058), we wondered if the growth in hyaline cartilage produced by MTOs would be more closely associated with the increased expression of articular cartilage gene markers but not others. Indeed, we only observed a significant increase in the expression of articular cartilage marker genes (GO:0061975; p=0.031) and not bronchus or trachea cartilage development (GO:0060532; GO:0060534).

**Table 2.** Change of gene expression in biological processes (GO) of interest

| **GO Terms** | **Description** | **Count in network<sup>1</sup>** | **p-value<sup>2</sup>** | **Significance of increase<sup>3</sup>** |
| --- | --- | --- | --- | --- |
| **GO:0061036** | **positive regulation of cartilage development** | **29/32** | **0.0053** | **\*\*** |
| **GO:0060591** | **chondroblast differentiation** | **4/5** | **0.0039** | **\*\*** |
| **GO:0060536** | **cartilage morphogenesis** | **8/11** | **0.019** | **\*** |
| GO:0003417 | growth plate cartilage development | 18/18 | 0.058 |  |
| GO:0061037 | negative regulation of cartilage development | 26/32 | 0.17 |  |
| GO:0032331 | negative regulation of chondrocyte differentiation | 21/25 | 0.17 |  |
| GO:0003413 | chondrocyte differentiation involved in endochondral bone morphogenesis | 15/15 | 0.24 |  |
| **GO:0061975** | **articular cartilage development** | **3/3** | **0.031** | **\*** |
| GO:0060532 | bronchus cartilage development | 2/2 | 0.8 |  |
| GO:0060534 | trachea cartilage development | 6/9 | 0.6 |  |

<sup>1</sup> number of genes expressed/genes associated with the GO term (retrieved by biomaRt v2.46.3)
<sup>2</sup> one-tailed Wilcoxon test comparing normalized and log-transformed transcript counts between week 8 and week 15
<sup>3</sup> \*, p-value<0.05; \*\*, p-value <0.01

To summarize, we observed that protein products of genes with decreased expression in MTOs from week 8 to week 15 were clustered around components of Wnt signaling pathways that promote ossification while those with increased expression formed prominent networks around TGF-beta/BMP signaling pathways. Moreover, the observed molecular signaling clusters were associated with a unique increase in the gene expression of articular cartilage development.

### Transcriptomic comparisons between MTOs and lower limb chondrocytes cross human life stages

Because MTO RNA-seq revealed a unique increase in articular cartilage development, we hypothesized that there would be a strong correlation between MTOs and human growth plate chondrocytes and/or articular chondrocytes in the expression of genes specific to chondrocytes^26^. We first reprocessed existing RNA-Seq data collected from human embryonic limb bud (week 6)^26^, growth plate chondrocytes (week 14, 15, 16, and 18)^26^, knee chondrocytes (week 17, adolescent, and adult)^27,28^, as well costal chondrocytes (~70-year-old adults)^6^. We validated 325 genes known to be specifically expressed by chondrocytes^26^; we conducted PCA on these genes and found that the effect of different studies is minimal as gene expression was clustered by life stages rather than particular studies (Fig. 4a). Specifically, fetal tissues (growth plate and knee chondrocytes) clustered together while cartilage tissues from adolescents, adults, and 70-year-olds (knee and costal chondrocytes)—collectively addressed as post-*in utero* tissues—located in close proximity to each other (Fig. 4a). We highlight that 15-week MTOs were most similar to 6-week human limb bud cartilaginous tissues (Fig. 4a). Then, we used the Pearson correlation coefficient to examine the correlation among all samples and genes included in the PCA analysis. Agreeing with previous GO analysis and PCA, while all MTOs showed >60% correlation with human chondrocytes and cartilage tissues from all life stages, 15-week MTOs showed an even stronger correlation (>76%) with 6-week human limb bud and 14-week and 15-week fetal growth plate chondrocytes (Fig. 4b; Table S9). Moreover, we found that MTOs, `on average for previously described genes of interest, showed a significantly higher (p<0.0001; 95% CI [0.053, +∞]) correlation with fetal chondrocytes than post`-*in utero* knee chondrocytes; the average correlation between MTOs and fetal tissues was 71% while that for post-*in utero* tissues was 65%.

**Fig. 4.**
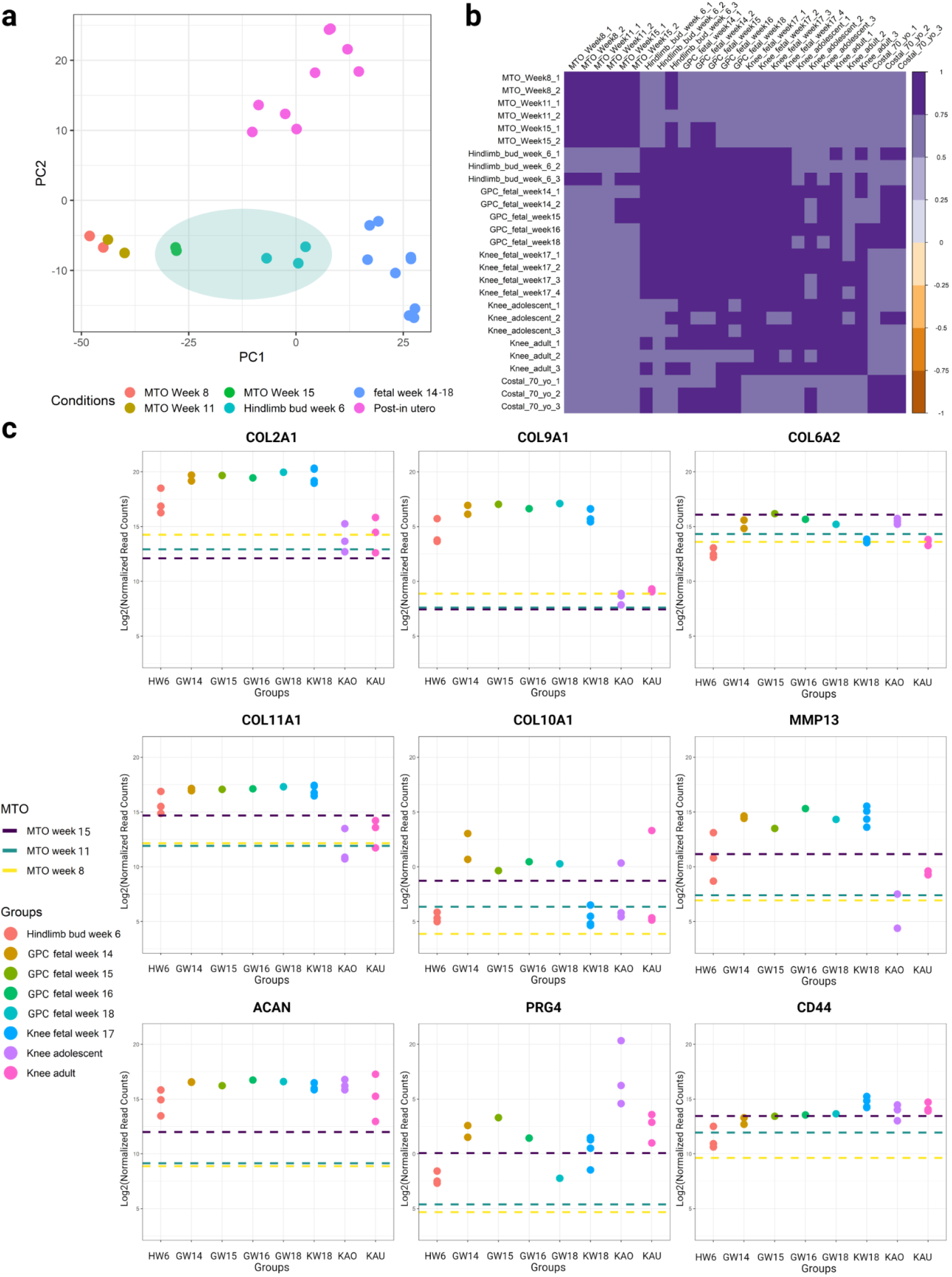
Transcription signatures in MTOs are comparable with human lower limb chondrocytes. **a.** Principal component analysis (PCA) on 325 chondrocyte-specific genes in MTOs and human lower limb chondrocytes. **b.** Pearson correlation plot of 325 chondrocyte-specific genes in MTOs and human lower limb chondrocytes. **c.** Representative comparisons of marker transcripts (*COL2A1, COL9A1, COL6A2, COL11A1, COL10A1, MMP13, ACAN, CD44, PGR4*) in MTOs and human lower limb chondrocytes. MTOs, multi-tissue organoids; GPC, growth plate chondrocytes.

We further examined the expression of known collagen genes (*COL2A1, COL9A1, COL6A2, COL11A1, COL10A1*)^43^, hypertrophic markers (*COL10A1, MMP13*), and some components reflecting the secretory functions of chondrocytes (*ACAN, CD44, PGR4*) in MTOs as compared to human lower limb chondrocytes. We found that transcripts of components for ECM were generally more abundant from *in utero* than post-*in utero* lower limb chondrocytes, while those for secretory molecules showed more variable trends. Although the expression of genes encoding for type 2/9 collagen (*COL2A1, COL9A1*) was lower in 15-week MTOs, the expression from 8- and 11-week MTOs fell in the range defined by examined human tissues. The expression of other major collagen components making up the non-hypertrophic region of cartilage (*COL6A2* and *COL11A1*) and hypertrophic markers (*COL10A1, MMP13*) were not noticeably different between MTOs and human lower limb cartilaginous tissues. Among MTOs collected at different time points, despite falling in expression ranges of human tissues, 15-week MTOs showed slightly lower collagen contents and higher hypertrophic gene expression. As for transcripts of secretory molecules, 15-week MTOs showed similar expression (*PGR4, CD44*) to human tissues except for lower *ACAN*. We note that the normal expression of *PRG4*, a gene encoding for a large proteoglycan synthesized by chondrocytes located at the surface of articular cartilage and by some synovial lining cells^44^, offers additional support to the articular developmental trajectory spontaneously taken by MTO chondrocytes. In short, we found that transcripts levels of genes specifically expressed in human growth plate chondrocytes were strongly correlated between 15-week MTOs and human lower limb cartilaginous tissues.

## Discussion

Chondrogenesis and hyaline cartilage production *in vitro* is needed to provide chondrocytes for cartilage regenerative therapies as well as for *ex vivo* OA modeling. Despite recent advances in hiPSC-based *in vitro* chondrogenesis^5,6^, the development of simple and scalable protocols to generate chondrocytes for therapeutic purposes as well as the understanding of dynamic cell behaviours in long-term cultures are yet to be accomplished. Here, we report the spontaneous emergence and robust growth of hyaline cartilage in hiPSC-derived MTOs grown in xeno- and feeder-free, chemically-defined cultures, making this protocol amenable for clinical good manufacturing practices (cGMP). Furthermore, the relative technical simplicity of the process also makes it suitable for robotic cell culture and scaled-up manufacturing. By characterizing and analyzing the transcriptome changes during the phenotypic transition of MTOs, we provide a mechanistic foundation for future organoid and related tissue engineering aiming at rapid cartilage production to thrive upon.

We first described the morphology and characteristic histochemical and immunohistochemical features of the MTO-cartilage. Specifically, prominent hyaline cartilage emerged by week 8 of culturing MTOs and continued to expand in size. Further, we found that COL2A1 IHC staining increased over time as the cartilage matured despite decreased transcript level, Type VI collagen was present especially in pericellular locations, while COL1A1 showed restricted immunoreactivity most evident at the periphery of cartilage at the junction with surrounding tissue, and COL10A1 was generally not detectable above background. We conclude that the hyaline cartilage produced by MTOs is most similar to articular hyaline cartilage. Key collagen and ECM compositions affect the functional performance of cartilages. Cartilage with greater tensile strength contains a higher content of collagen type II/IX, aggrecan, and a lower content of collagens type I/X^45-47^. Higher aggrecan content in the ECM allows more capacity for the cartilage to withstand compression^45-47^. Our findings, therefore, suggest that MTO cartilage will have biomechanical properties similar to articular cartilage including resistance to repetitive forces of compression and shear.

MTOs were formed by the previously reported CO^25^ undergoing a substantial and self-organized transition without interventions beyond the change of flask and regular media change. In contrast to the increase in neural cells in hiPSC-derived chondrocyte populations observed by Wu et al^6^, we observed a consistent decrease in neural lineages in MTOs collected at different time points. It is unclear whether cells of neural lineages during MTO development underwent senescence or fate switching. Regardless, neural crest cells (NCC) may have emerged from neural tube-like structures present in CO, which later spontaneously formed mesodermal derivatives, such as chondrocytes. Induction of chondrocytes from hiPSC through NCC intermediates has been recently reported by two independent groups^5,7^. Moreover, RNA-seq of MTOs agreed with the phenotypic transition of MTOs from week 8 to week 15, where we observed favored BMP pathway signaling and increased mesodermal processes associated with cartilage production. Interestingly, MTOs showed spontaneous increases in the expression of all genes coding for previously reported inductive factors essential for chondrogenesis, including *BMP2, BMP6, FGF2, GDF5*, and *TGFB1*^5, 8^; this suggests that MTO chondrocytes likely shared similar, albeit spontaneous, developmental trajectories with other hiPSC-derived chondrocytes^5,8^. Together, although our data only demonstrated correlative relationships, existing literature surrounding chondrogenic differentiation and the conserved roles of BMP and FGF signaling pathways suggests that favored expression of components in BMP signaling pathways resulted in the increased cartilage production in MTOs. This was further supported by the prominent clustering of TGF-beta/BMP signaling pathways formed by gene products encoded by genes with increased expression during cartilage development in MTOs. Furthermore, we observed a probable downregulation of components in the Wnt signaling pathways related to cartilage development. This is in agreement with previous reports indicating that Wnt signaling pathways inhibit chondrogenesis and/or promote ossification^39,48,49^. With regard to hiPSC-derived chondrocytes, Wnt signaling was recently shown to cause off-target differentiation during chondrogenesis by Wu et al^6^. We were not surprised to see the overlaps of ossification and TGF-beta/BMP pathways because markers for cartilage development—such as *BMP2* and *BMP6*—are largely expressed throughout the hypertrophic change of chondrocytes^34^. Therefore, the increase in TGF-beta/BMP pathways involved in ossification should not be solely interpreted as progress towards ossification. This was further supported by the non-significant change in expression of genes involved in chondrocyte differentiation associated with endochondral bone morphogenesis (GO:0003413). To summarize, it is likely that 1) the favoring of BMP signaling pathways over neural FGF pathways promoted mesoderm formation and 2) the combination of upregulated TGF-beta/BMP signaling and downregulated Wnt signaling components resulted in the long-term maintenance of cartilage growth and its specific increase expression of articular cartilage development marker genes. Our findings did not, however, rule out the involvement of other pathways or the differential signaling within a subset of pathways. For example, the crosstalk between canonical and noncanonical TGF-beta signaling pathways may also contribute to the maintenance of cartilage^50^ in MTOs. Extensive research to understand and manipulate the precise spatiotemporal expression of aforementioned components during MTO development, specifically in chondrocytes and their precursors, would be needed to establish definitive causative effects. MTOs may serve as human-specific models for disease modeling and drug testing due to the developmental dynamics of key chondrogenic pathways.

To evaluate the similarities and differences between cartilage^51^ from MTOs and human tissues, we reprocessed existing RNA-Seq data obtained from human fetal limb bud, and chondrocytes from fetal growth plate and knee, and adolescent and adult chondrocytes. We found that the 15-week MTOs showed a strong correlation (>76%) with human fetal limb bud and growth plate chondrocytes for growth plate chondrocyte-specific gene expression. Together with comparable levels of *PRG4* transcripts, the result suggests that MTOs spontaneously adopt the trajectory of articular cartilage development during long-term culture. Although there is a slight difference in the expression of type II/IX collagen transcripts, the expression of other cartilage components and secretory chondrocyte molecules were similar. The expression of collagen contents in MTOs may change and/or improve as the environment becomes more physiological. Long-term implantation of chondrocytes in large animals may shed light on the therapeutic value of MTO-derived chondrocytes.

To conclude, the long-term culture of hiPSC-derived MTOs results in the spontaneous emergence of mesoderm-derived articular cartilaginous tissues and the MTO cartilage resembles fetal limb bud and growth plate chondrocytes. Despite previous reports of hyaline cartilage appearance facilitated by chondrogenic induction of hiPSC *in vitro*^5-8^, MTO-generated cartilage stands out as they are larger, more mature, and can be maintained in long-term cultures. This process was also self-organized with a comparably simpler, xeno- and feeder-free, 3D culturing protocol, making production easily adaptable to cGMP production and amenable to scaled-up commercial manufacturing. The transition from CO ectodermal trajectory to the commitment of articular cartilage development in MTOs reported here further illustrates the remarkable self-organization capability of organoids.

## Methods

### Generating hiPSC-derived multi-tissue organoids (MTOs)

iPSC lines referred to as 1024 (ATCC-BYS0110, Cat. #ACS-1024) and 9-1 (UMN PCBC16iPS/vShiPS 9-1)^25^ were expanded in culture on vitronectin (VTN-N, Thermofisher Scientific, Cat. #A14700) in Essential 8 Medium (E8, Thermo Fisher Scientific Life Sciences). iPSCs were harvested using sodium citrate buffer, briefly centrifuged, and MTO induction initiated by resuspension in 40 μL of the fluid of hydration from Cell-Mate3D μGel 40 Kit (BRTI Life Sciences, Two Harbors, MN, USA) which was then transferred to one well of a 6-well ultra-low attachment plate (Costar™ Ultra-Low Attachment Microplates, Corning Life Sciences, Corning, NY, USA) containing 5 ml of E8 medium and incubated for 24 hrs. Cells and culture medium were then transferred to a G-Rex 100 bioreactor (Wilson Wolf, New Brighton, MN) and incubated at 37°C in 5% CO_2_. E8 culture medium containing 1% antibiotic-antimycotic (Gibco Thermo Fisher Scientific) was changed every 3-4 days over the entirety of MTO culture.

### Histology and Immunohistochemistry

MTOs were harvested on weeks 8 and 11 (both cell lines) and week 30 (1024 only), placed in 10% neutral buffered formalin solution and fixed at room temperature for 3.5 hours. After fixation, samples were transferred to 70% ethanol solution until they were processed for routine paraffin embedding. Samples were then sectioned 4-μm thick, deparaffinized, rehydrated, and routinely stained with hematoxylin and eosin (H&E) and Alcian blue. For immunohistochemical staining, sections were cut at 4 μm, deparaffinized, and rehydrated, followed by incubation with 3% hydrogen peroxide to quench endogenous peroxidase activity and 15 minutes in serum-free protein block (DAKO, Glostrup, Denmark). Sections were then subjected to appropriate antigen retrieval methods (if needed) and incubated with the primary antibody at room temperature for 60 minutes (Table S10). Color development was done using EnVision FLEX DAB+ substrate chromogen system (Cat.# GV825, Agilent-Dako, Santa Clara, CA, USA). Stained sections were examined with an Olympus BH-2 microscope (Olympus America, Center Valley, PA) and imaged with a SPOT Insight 4 megasample digital camera and SPOT Advanced software (Diagnostic Instruments Inc., Sterling Heights, MI).

### Morphometry

Aggrecan IHC stained histologic sections of MTO biological replicates at 8 weeks (n=7) and 11 weeks (n=5) were analyzed for aggrecan staining area fraction (staining area/total area of tissue) using a Nikon Eclipse E-800M bright field/fluorescence/dark field microscope equipped with a Nikon DXM1200 high-resolution digital camera. Images for histomorphometry were analyzed using ImageJ2/Fiji software (National Institutes of Health, open-source). Values are reported as area fraction (%) ± standard deviation.

### RNA-seq of MTOs and data generation

MTOs were disaggregated into single cell or small clusters to enhance RNA extraction. MTOs were pipetted from the bioreactor, rinsed in PBS and suspended in 2 ml trypsin (0.025%, Sigma Aldrich) for 2 minutes at 37°C. Next, a scalpel was used to mechanically disrupt the MTOs in a conical tube in 600 μg DNAse1 in an additional 3 ml of 0.025% trypsin and this was incubated for 5 minutes at 37°C. MTOs were then dissociated by a stream of cold Hank’s Balanced Salt Solution (HBSS; Life Technologies) over a 100 μm filter (BD Biosciences, San Jose, CA). Organoid tissue retained on the filter was then back-flushed with 25 ml of cold HBSS and this tissue was further disaggregated in 2 ml Collagenase II (Celase GMP, Worthington Biochemical Corporation, Lakewood, NJ) also containing 200 μg DNAse1, for 5 minutes at 37°C. After incubation, tissue was gently triturated with a 5 ml pipet and an additional 3 ml of Collagenase II with 300 μg DNAse1 added, and this preparation was incubated for 5 minutes at 37°C. The cells were then centrifuged at 350 x g for 3 minutes at 4°C, the cell pellet was resuspended in cold HBSS, centrifuged again and resuspended in HBSS.

The MTO cell preparations were then lysed in RLT buffer (Qiagen, Hilden, Germany) and RNA isolated from cell lysates using the RNeasy Plus mini kit (Cat. No. / ID: 74134Qiagen, Hilden, Germany) according to the manufacturer’s instructions. Extracted RNA was then quantified by RiboGreen RNA assay (Thermo Fisher Scientific, Waltham, MA) and quality/size analyzed by Agilent BioAnalyzer (Agilent Technologies, Santa Clara, CA). 2 x 50bp FastQ paired-end reads for 6 samples (n=62.4 Million average per sample) were trimmed using Trimmomatic (v0.33) enabled with the optional “-q” option; 3bp sliding-window trimming from 3’ end requiring minimum Q30. Quality control on raw sequence data for each sample was performed with FastQC. Read mapping was performed via Hisat2 (v2.1.0) using the Human genome (GRCh38) as reference. Gene quantification was done via Feature Counts for raw read counts. Existing RNA-seq data were processed using the same pipeline.

### RNA-seq data analysis

Raw read counts (CPM) were used as input for differentially expressed genes (DGE) analysis by DESeq2 package (v1.30.1) in R (v4.0.5)^52^. P-values were adjusted (p-adj) using Benjamini-Hochberg correction. The significant term was determined by using a cut-off of 0.05 (FDR corrected p<0.05) and minimum 2x Absolute Fold Change. TopGo package (v2.42.0)^53^ was used to carry out ontology enrichment of DGE results^35^. Enrichment results were visualized using ClusterProfiler (v3.18.1)^54^. Cut-offs for p-value (after applying the Benjamini-Hochberg correction) and q-value were 0.05 and 1, respectively. Genes with expression FDR<0.05 were selected and uploaded to STRING (v11) was used for functional protein association networks for probable gene products; Euclidean distances were used to cluster gene products^55^. Local network clusters were downloaded from STRING analysis^55^.

### Additional statistical information

The one-tailed Welch’s t-test was used to compare histomorphometric measurements on MTO histologic sections. P-value was reported (α=0.05). Two biological replicates for MTOs were used for RNA-Seq. All gene expression data used for visualization and statistical tests were first normalized and transformed using rlog() and assay() in DESeq2^52^. The one-tailed Wilcoxon signed-rank test was used to acquire statistical results for the change in expression of grouped genes; p-values were reported (α=0.05). The one-tailed pairwise t-test was used to compare the expression of single genes; p-values were reported (α=0.05). Confidence intervals were reported when applicable.

## Supporting information

TableS1_15vs8_DGE.csv

TableS2_MarkerGenes.csv

TableS3_decreased_enrichment.tsv

TableS4_increase_enrichment.tsv

TableS5_enrichment.NetworkNeighborAL_decreased.tsv

TableS6_enrichment.NetworkNeighborAL_increased.tsv

TableS7_ExpressionTable_for_Table2.xlsx

TableS8_ExpressionofChondrocyteSpecificGenes.csv

TableS9_Pearson_Correlaiton_Table.csv

## Data Availability

All RNA-seq data and raw read counts generated for the analyses herein are publically available within the NCBI SRA archive under project number: GSE184007.

## Acknowledgements

The authors would like to acknowledge the University of Minnesota Masonic Cancer Center Comparative Pathology Shared Resource for excellent technical support by Paula Overn in the performance of all histology sample preparation and performance of immunohistochemical staining, and Dr. Katalin Kovacs for performing the histomorphometric analysis of the MTOs. The University of Minnesota Genomics Center kindly assisted by generating the Illumina sequencing data reported herein. The authors acknowledge the Minnesota Supercomputing Institute (MSI) at the University of Minnesota for providing resources that contributed to the research results reported within this paper. Figures were created or organized using BioRender (biorender.com).

Research reported in this publication was supported by the National Center for Advancing Translational Sciences of the National Institutes of Health Award Numbers grant UL1TR002494 and UL1TR002377. The content is solely the responsibility of the authors and does not necessarily represent the official views of the National Institutes of Health. Additional support was provided from the Minnesota Partnership for Biotechnology and Medical Genomics through the Translational Product Development Fund (TPDF). The study was also supported by internal University of Minnesota grants from the Faculty Research Development Grants fund, the College of Veterinary Medicine Comparative Medicine Signature Grant Program, and the Institute for Engineering in Medicine collaborative grants program.

## Supplementary Information

Supplementary Information: Supplementary information

Supplementary Table 1: TableS1_15vs8_DGE.csv

Supplementary Table 2: TableS2_MarkerGenes.csv

Supplementary Table 3: TableS3_decreased_enrichment.tsv

Supplementary Table 4: TableS4_increase_enrichment.tsv

Supplementary Table 5: TableS5_enrichment.NetworkNeighborAL_decreased.tsv

Supplementary Table 6: TableS6_enrichment.NetworkNeighborAL_increased.tsv

Supplementary Table 7: TableS7_ExpressionTable_for_Table2.xlsx

Supplementary Table 8: TableS8_ExpressionofChondrocyteSpecificGenes.csv

Supplementary Table 9: TableS9_Pearson_Correlaiton_Table.csv

