## Supplementary Information for "Self-organized emergence of hyaline cartilage in hiPSC-derived multi-tissue organoids"

### Author Information

Manci Li[1,2], Juan E. Abrahante[3], Amanda Vegoe[4,5,6], Yi Wen Chai[4,5,6], Beth Lindborg[4,5,6], Ferenc Toth[7], Peter A. Larsen[1,2], Timothy D. O'Brien[4,5,6]

[1]Department of Veterinary and Biomedical Sciences, University of Minnesota, St. Paul, MN 55108

[2]Minnesota Center for Prion Research and Outreach, College of Veterinary Medicine, St. Paul, University of Minnesota, MN 55108

[3]University of Minnesota Informatics Institute, University of Minnesota, Minneapolis, MN 55455

[4]Stem Cell Institute, University of Minnesota, Minneapolis, MN 55455

[5]Department of Veterinary Population Medicine, University of Minnesota, St. Paul, MN 55108

[6]Sarcio, Inc., Minneapolis, MN 55455

[7]Department of Veterinary Clinical Sciences, University of Minnesota, St. Paul, MN 55108

**Fig. S1**

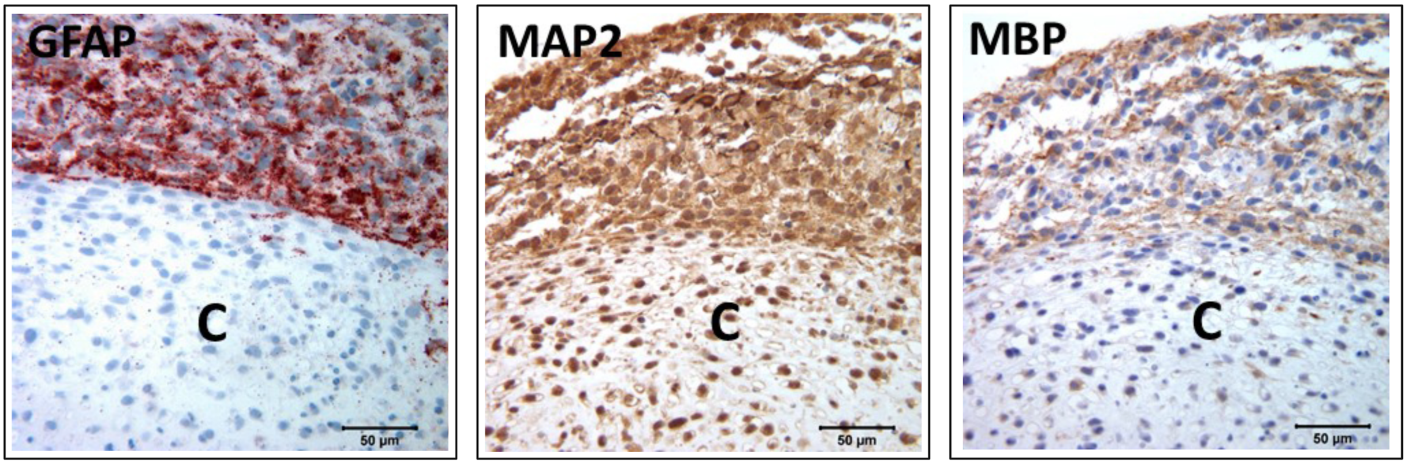


Immunohistochemical staining for brain cell lineages in 1024 MTOs at 8 weeks demonstrates strong labeling for glial fibrillary acidic protein (GFAP, astrocytes), microtubule-associated protein 2 (MAP2, neurons), and myelin basic protein (MBP, oligodendrocytes) in surface regions of the MTO adjacent to the chondrogenic region (C).

**Fig. S2**

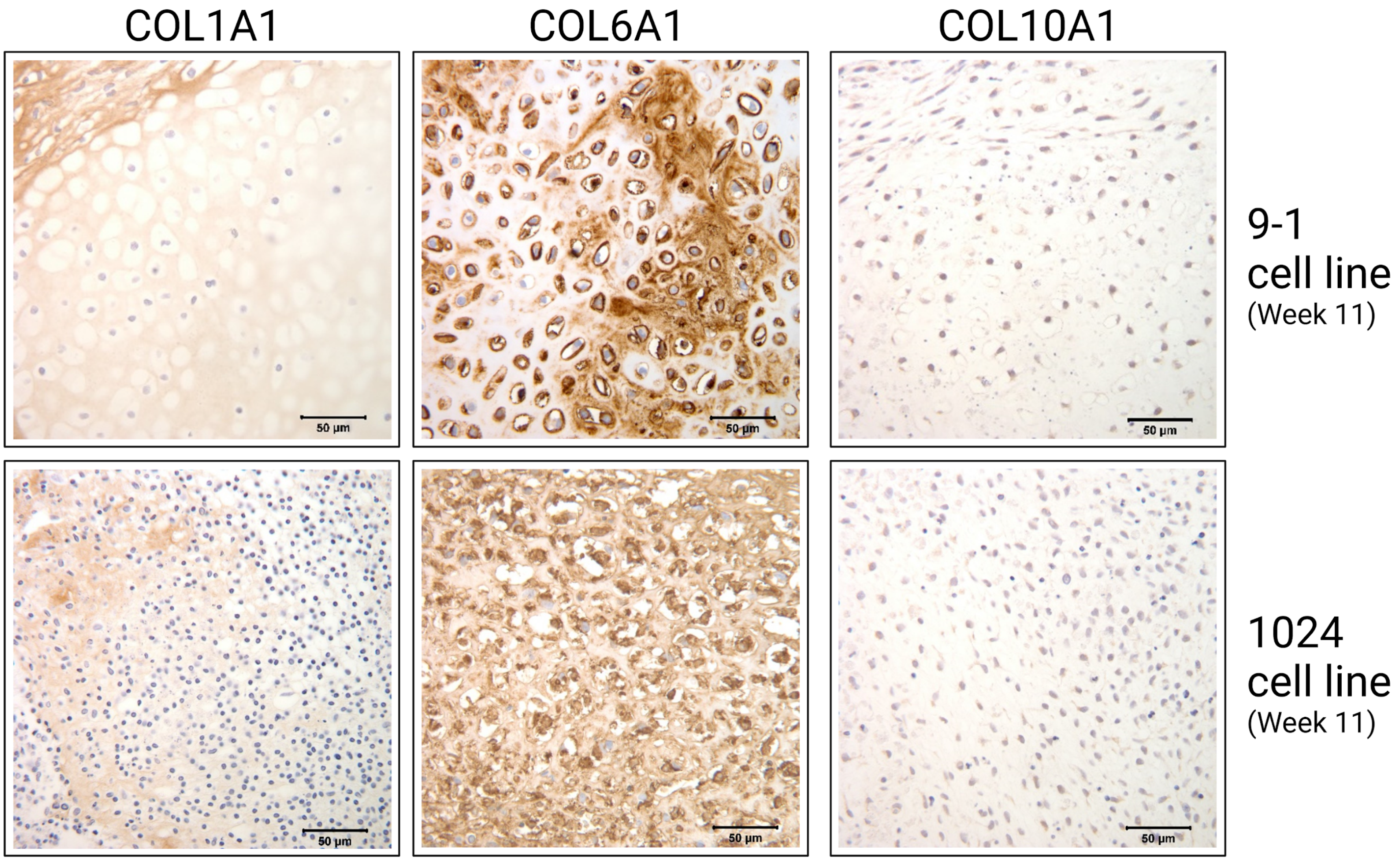


11-week MTOs from 9-1[1] and 1024 cell lines showing some peripheral weak to moderate staining for type I collagen (COL1A1), pericellular and some patchy matrix staining for type VI collagen (COL6A1), and negative staining for type X collagen (COL10A1).

**Fig. S3.**

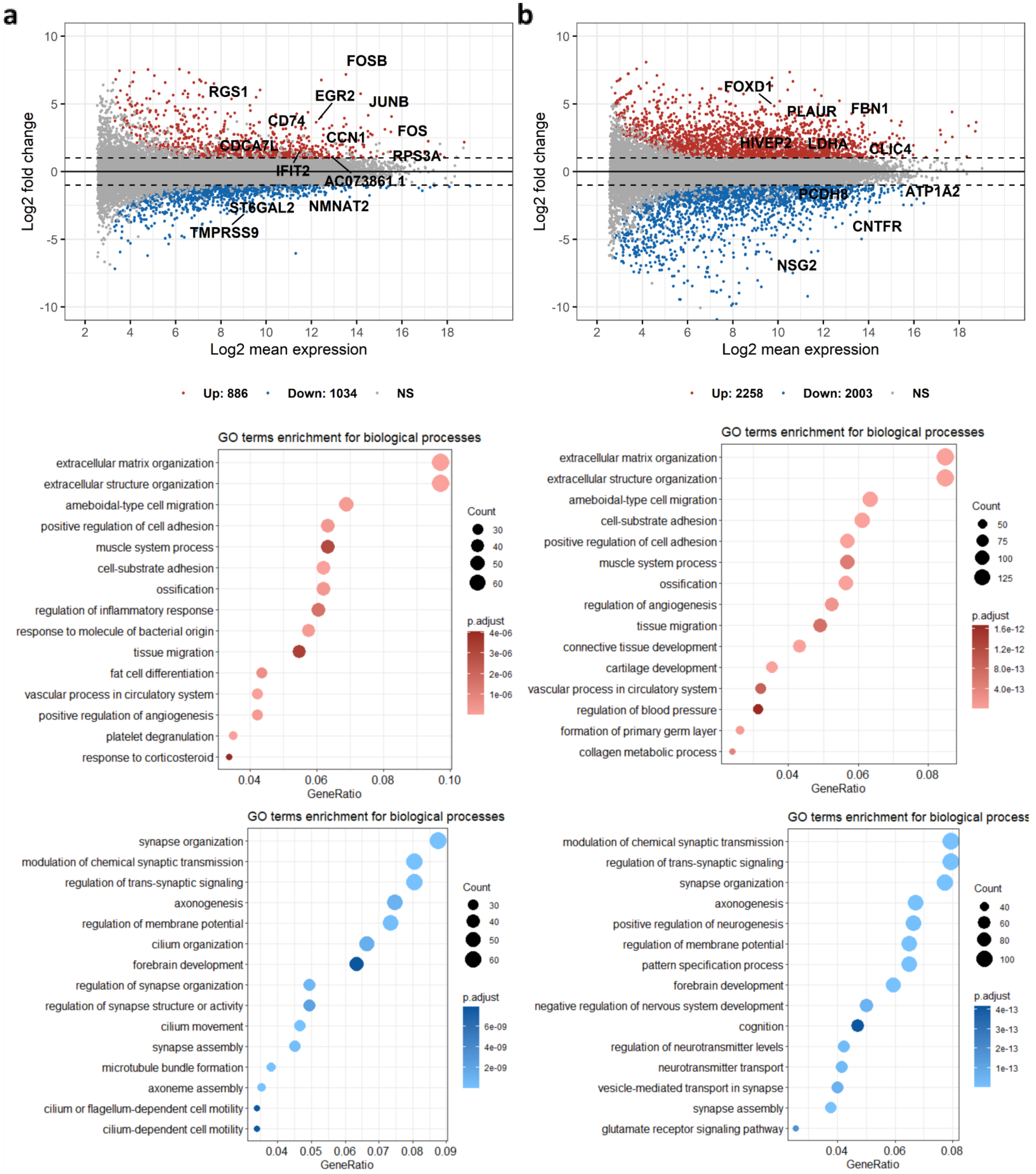


Differential gene expression and gene ontology enrichment analyses (week 11 vs week 8 and week 15 vs week 11). **a.** Differential gene expression, gene ontology enrichment for upregulated and genes, gene ontology enrichment for downregulated genes between week 11 and week 8. **b.** Differential gene expression, gene ontology enrichment for upregulated and genes, gene ontology enrichment for downregulated genes between week 15 and week 11.

**Fig S4.**

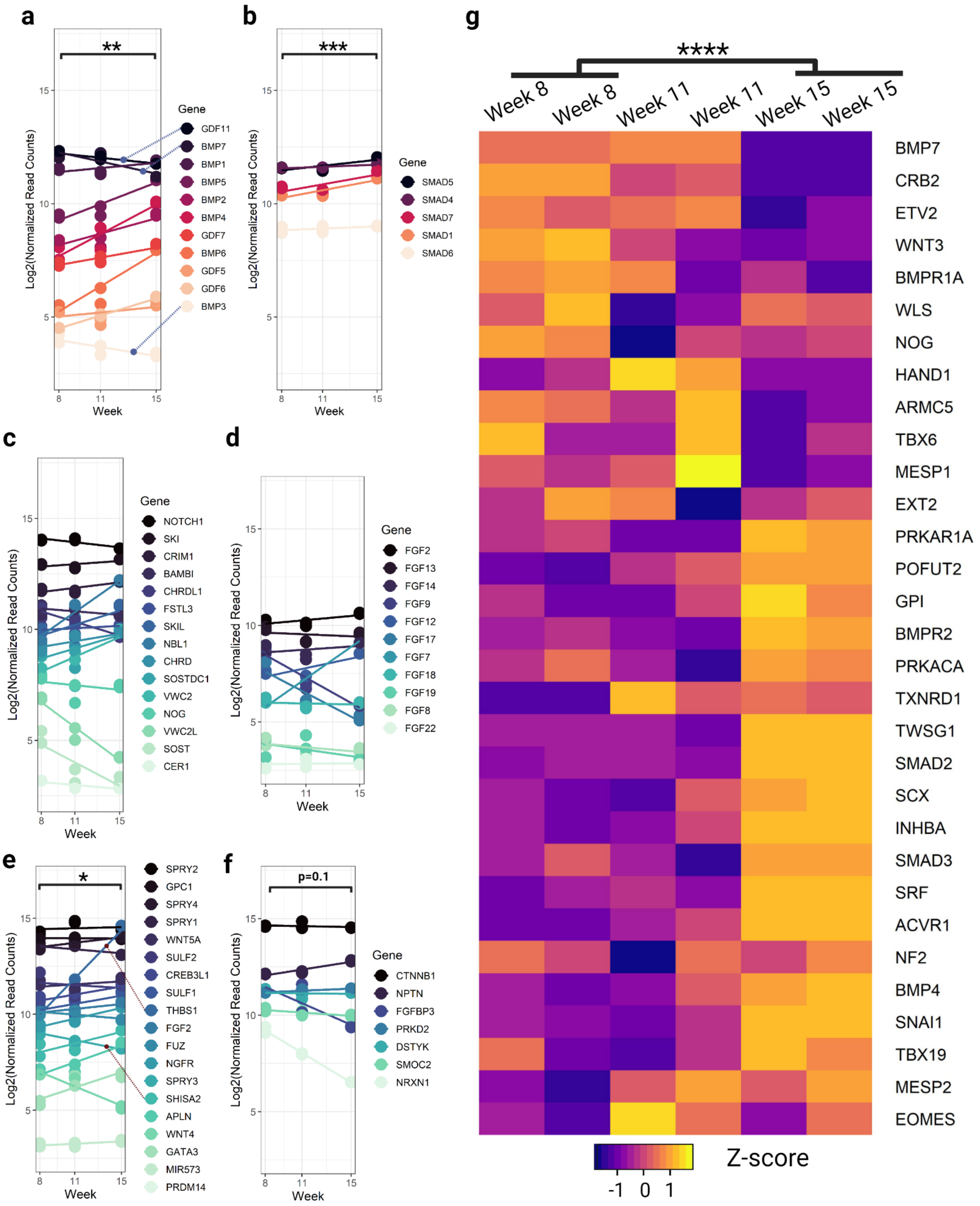


Gene expression plots of gene lists of interests. **a.** Bone morphogenic proteins (BMP). **b.** SMAD proteins. **c.** BMP antagonists. d. Neural FGFs. **e.** Negative regulation of FGFR pathway (GO:0040037). **f.** Positive regulation of FGFR pathway (GO:0045743). **g.** Mesoderm formation (GO:0001707).

**Table S9. Antibodies used for histology and immunohistochemistry.**

| Antibody | Type of Antibody | Manufacturer/Cat. No. | Dilution | Antigen Retrieval |
|---|---|---|---|---|
| Aggrecan (ACAN) | Goat polyclonal | R&D Systems, Cat# AF1220 | 1:200 | None |
| Type I Collagen (COL1A1) | Mouse monoclonal | Sigma c2456 | 1:100 | Proteinase K |
| Type II Collagen (COL2A1) | Mouse monoclonal | Developmental Studies Hybridoma Bank CIIC1 | 1:10 | Pepsin |
| Type VI Collagen (COL6A1) | Rabbit polyclonal | abcam ab6588 | 1:400 | Citrate |
| Type X Collagen (COL10A1) | Rabbit polyclonal | abcam ab58632 | 1:400 | Typsin |
| Glial Fibrillary Acidic Protein (GFAP) | Rabbit polyclonal | Dako Cat.#Z0334 | 1:1000 | None |
| Microtubule Associated Protein 2 (MAP2) | Chicken polyclonal | abcam ab5392 | 1:4000 | Citrate |
| Myelin Basic Protein (MBP) | Rabbit polyclonal | abcam ab40390 | 1:3000 | Citrate |
